# Phylogenetic clustering of microbial communities as a biomarker for chemical pollution

**DOI:** 10.1101/2025.02.12.637817

**Authors:** Thomas P. Smith, Rachel Hope, Thomas Bell

## Abstract

Microbial communities play a critical role in ecosystem functioning and offer promising potential as bioindicators of chemical pollution in aquatic environments. Here we examine the responses of both bacterial isolates and microbial communities to a range of pollutants, focusing on the phylogenetic predictability of their responses. We found that bacterial isolates exhibited a strong phylogenetic signal in their growth responses, with closely related taxa responding similarly to chemical stress. In microbial communities, pollutants that significantly impacted isolates also reduced community diversity and growth, causing shifts in community structure toward increased phylogenetic clustering, suggesting environmental filtering. The net relatedness index (NRI) effectively captured these shifts, indicating its potential as a simple metric for monitoring pollution. Our findings highlight the predictability of microbial responses to pollution and suggest that microbial-based bioindicators, coupled with rapid sequencing technologies, could transform environmental monitoring.

## Introduction

Freshwater ecosystems are functionally diverse environments which play a crucial role in maintaining global biogeochemical cycles, regulating the flow and transformation of key nutrients and greenhouse gases (Bani et al., 2022; Sagova-Mareckova et al., 2021). However, these natural environments are coming under increasing pressure from anthropogenic stressors, such as toxic chemical pollutants (Stehle & Schulz, 2015; Thompson et al., 2016). Microbes are central to the ecological and biogeochemical processes occurring within freshwater environments, driving biogeochemical cycles and providing key ecosystem services (Grossart et al., 2020; Sagova-Mareckova et al., 2021). Microbes are particularly responsive to environmental stressors due to their high metabolic rates and rapid growth (Sagova-Mareckova et al., 2021). Microbes therefore have the potential to be used as rapid ‘biosensors’ that detect and track chemicals of concern (Smith et al., 2024), particularly because many chemical pollutants show off-target impacts on microbial populations (Lindell et al., 2024).

The responses of whole communities of microbes to environmental perturbations, can be observed through changes in their composition and structure (Astudillo-García et al., 2019). Prior studies of microbial communities have identified individual taxa that can be used as biomarkers for specific contaminants (Sagova-Mareckova et al., 2021; Wijaya et al., 2023). In addition, “keystone” or “multitask” taxa have been identified that are hallmarks of multiple environmental conditions and predict general environmental pollution or quality (Sperlea et al., 2021; Wang et al., 2020). A key limitation of indicator taxa is that there are many factors that determine whether a specific taxon may or may not be present in a given environment, and many of those factors are unrelated to environmental stress.

An alternative to indices based on specific taxa is to use taxonomy-free properties of the community for biomonitoring (Cook et al., 2024). One proposed taxonomy-free way to measure disturbed ecosystems is through phylogenetic structure (Helmus et al., 2010). If habitat filtering is the dominant assembly process in disturbed ecosystems, closely related species that share similar traits will be selected, resulting in a loss of genetic diversity (Gainsbury & Colli, 2019; Helmus et al., 2010). We can extend this to the effects of chemical pollutants on microbes by considering how species would be filtered by chemicals with phylogenetically constrained effects (Fig. 1). If chemicals show fitness-limiting effects on similar microbes, in a phylogenetically constrained pattern, then we would expect to observe phylogenetic clustering in communities exposed to these chemicals (Fig. 1a). That is, if clades of closely related species are impacted and lost from the community, then the remaining, unimpacted species will also be more closely related than expected (Cooper et al., 2008; Webb et al., 2002). Simulations have shown that microbial community responses to global change are likely to be phylogenetically conserved (Amend et al., 2016), however it is unclear if microbial responses to chemical pollutants are phylogenetically conserved (Smith et al., 2024). If this is the case, microbial phylogenetic community structure may offer a simple metric to diagnose polluted or perturbed ecosystems.

**Figure 1.**
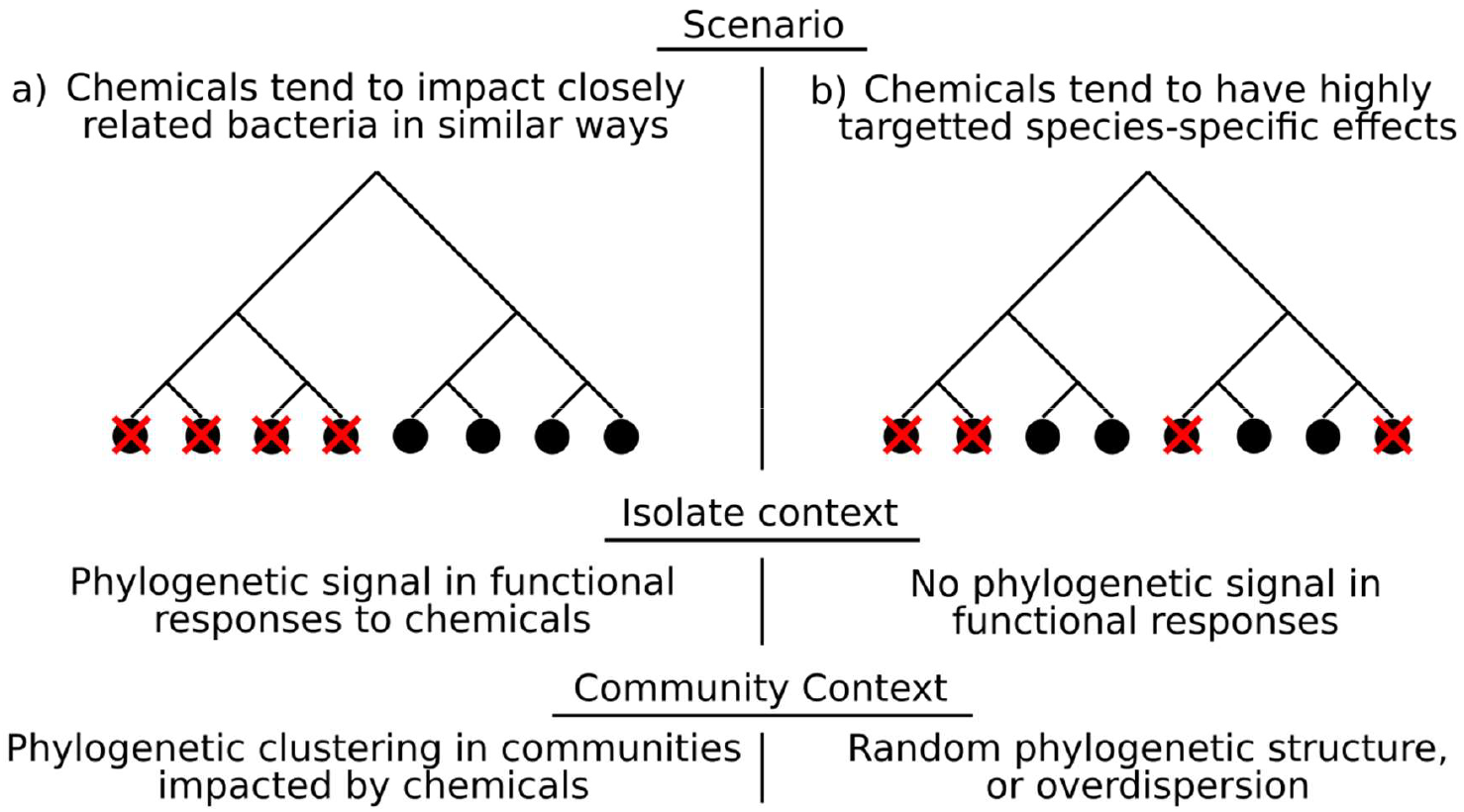
Phylogenetic structure in community responses to chemicals. **a**. If chemical stressors negatively affect microbes (indicated by a red cross) in a phylogenetically conserved way, i.e. similar taxa are all impacted by a given chemical, then we would expect to observe phylogenetic clustering in communities impacted by the same chemical as whole clades of taxa are filtered out. **b**. Conversely, if there is no phylogenetic component to which species are impacted by a chemical, we would not expect to see phylogenetic clustering when compared to community structure in the absence of a chemical stressor.

Recent work has shown that microbial community assembly is highly predictable and repeatable, resulting in set community structures (Pascual-Garcia et al., 2023). While communities may differ in their composition from site to site due to environmental differences or other factors (e.g. priority effects), we would nonetheless expect that phylogenetically related taxa will respond to pollutants in a structured, repeatable way, driven by their shared evolutionary history. Here we compared the responses of single species and whole microbial communities to a suite of chemical pollutants to investigate their potential for bioindicator use based on phylogenetic relatedness. We tested: (1) whether there was phylogenetic signal in the responses of bacterial isolates to chemical pollutants; (2) whether chemical pollutants altered the composition and phylogenetic structure of microbial communities; and (3) whether the responses of monocultured taxa are reflected in the community context.

## Results

We assayed the impact of 168 chemical pollutants on 26 strains of bacteria in our culture library, isolated from two different habitats. These habitats were stream sediments in Iceland, see (Smith et al., 2024), and soils from Silwood Park, Berkshire, see (Mombrikotb et al., 2022). The chemical pollutants were chosen based on a previous analysis of off-target effects of industrial and agricultural chemicals on human gut bacteria (Lindell et al., 2024). We quantified the growth of bacterial isolates by measuring the optical density of liquid cultures every hour for 72 hours (3 days) using an automated plate reader. We quantified the area under the growth curve (AUC) as an overall metric of growth. We calculated the ratio of AUC in the presence of a chemical pollutant to the AUC in the presence of DMSO (control) to find the relative growth (dAUC) in the presence of a chemical. Of the potential 4368 bacteria-chemical responses (26 strains X 168 chemicals), 242 were significantly different from the controls, i.e. the chemical tested had a significant impact on growth of the isolate (Dunnett’s test, Supplementary Figure S1). These significant responses involved only 41 of the 168 chemicals tested. Of these significant responses, most were negative (232/242, 95.9% of significant bacteria-chemical responses dAUC < 1), i.e. growth was significantly reduced in the presence of the chemical (Fig. 2a). The responses also displayed a nested pattern. That is, some more susceptible isolates were impacted by many chemicals, whereas some more resilient isolates were impacted by only a few chemicals, but these few were a subset of those impacting the susceptible isolates (Fig. 2a).

**Figure 2.**
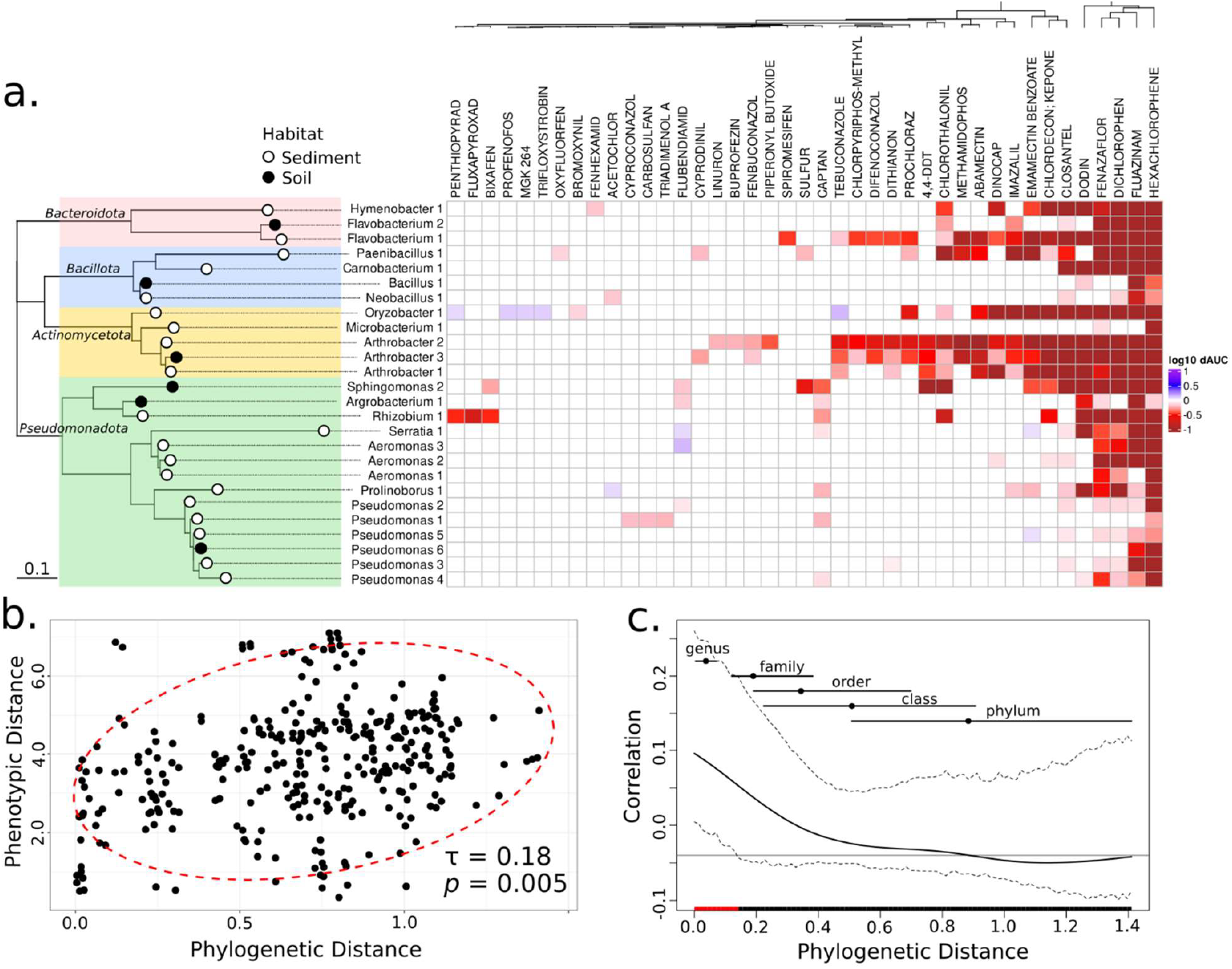
Phylogenetic signal in isolate responses to chemical pollutants. **a**. Heatmap of chemical impacts on isolate growth. Colours show the impact on growth on a log_10_ scale, i.e. -1 is 10x less growth, 0 is the same as control growth, +1 is 10x more growth. Dendrogram above shows hierarchical clustering of the chemical responses based on Euclidean distance. Rows are ordered by the phylogeny of the isolates built from their 16S sequences, shown to the left. Different phyla are highlighted by colour, scale bar shows phylogenetic distance. Only the chemicals with significant impacts on the growth of at least one isolate are shown. **b**. Pairwise phylogenetic distance plotted against pairwise phenotypic distance (from hierarchical clustering), with the Mantel test statistic (Kendall’s τ) used to compare the matrices. Red ellipse shows the 95% prediction region for the data distribution. **c**. Phylogenetic correlogram, representing the phylogenetic distances at which the correlation between traits is observed. Solid black line shows the Moran’s I index and dashed lines show the 95% confidence envelope. The horizontal black line indicates the expected value of Moran’s I with no phylogenetic correlation (null hypothesis). Red highlight indicates the range of phylogenetic distances with significant trait correlation (i.e. where the null hypothesis value of Moran’s I is outside of the 95% confidence interval). Points and horizontal lines represent the mean and range of phylogenetic distances between taxa in different levels of taxonomic organisation, respectively. For genus this is the mean pairwise distance between taxa of different genera within the same family, for family this is the mean pairwise distance between taxa of different families within the same order, and so on.

We assayed bacterial isolates from a wide range of phylogenetic groups, covering 4 bacterial phyla, obtained from stream sediments or soils (Fig. 2a). Similar taxa clustered together in their responses to the chemicals but were not perfectly grouped by taxonomy (Supplementary Figure S1). We tested for phylogenetic signal in the responses of the isolates using a Mantel test (using Kendall’s τ) comparing the pair-wise distance matrix computed from the chemical responses to the distance matrix of the phylogeny (Fig. 2b). We found a significant correlation between the phenotypic distance (responses to the chemicals) and the phylogenetic distance (Mantel statistic τ = 0.1829, *p* = 0.0054). This means that related bacteria tend to respond to the chemical treatments in the same way, despite being obtained from two disparate environments. We located where this phylogenetic signal occurred within the taxonomy, using a phylogenetic correlogram (Fig. 2c). A phylogenetic correlogram shows the correlation of a trait across species as a function of their phylogenetic distance, illustrating how trait similarity changes with increasing evolutionary separation (Keck et al., 2016).We find that the correlation occurs only at relatively short phylogenetic distances (<0.14), which at most equates to the distance between taxa in different genera or different families in our dataset (Fig 2c.).

We also assayed the growth responses of six whole bacterial communities obtained from the same stream sediments that the sediment isolates were cultured from to the same set of chemical stressors. The communities were generally resilient in their growth responses, clustering together and with the more resilient *Pseudomonas* isolates (Supplementary Figure S1). Only 10 of the chemicals had impacts on the overall growth of any of the communities, and none of the chemicals completely prevented community growth (Supplementary Figure S1). To understand how the community growth responses related to differences in community composition, we sequenced one of the communities (“community 9”) after growth in 23 of the chemicals, and the DMSO control. We selected a range of chemicals (see Supplementary Table S1), including those most impacting the communities’ growth, to understand whether impacts on growth were due to changes in community composition. We also selected some chemicals which elicited no significant impact on growth, to investigate functional redundancy - whether communities may have lost taxa but showed no change in functional response. We found remarkable consistency in the abundance profiles between replicates impacted by the same chemicals (Fig. 3a, Supplementary Figure S2). We also observed that the greatest changes in abundance profiles were in the presence of chemicals which also caused a change in community growth (Fig. 3a). These responses were very similar when aggregated to the genus-level, and exemplified by a loss of the *Aeromonas* taxa, with the exception of emamectin benzoate and methamidophos (Fig. 3a). However, we note that at the species-level, distinct taxonomic profiles arise in response to each of the chemicals which induce a growth response (Supplementary Figures S2 and S3).

**Figure 3.**
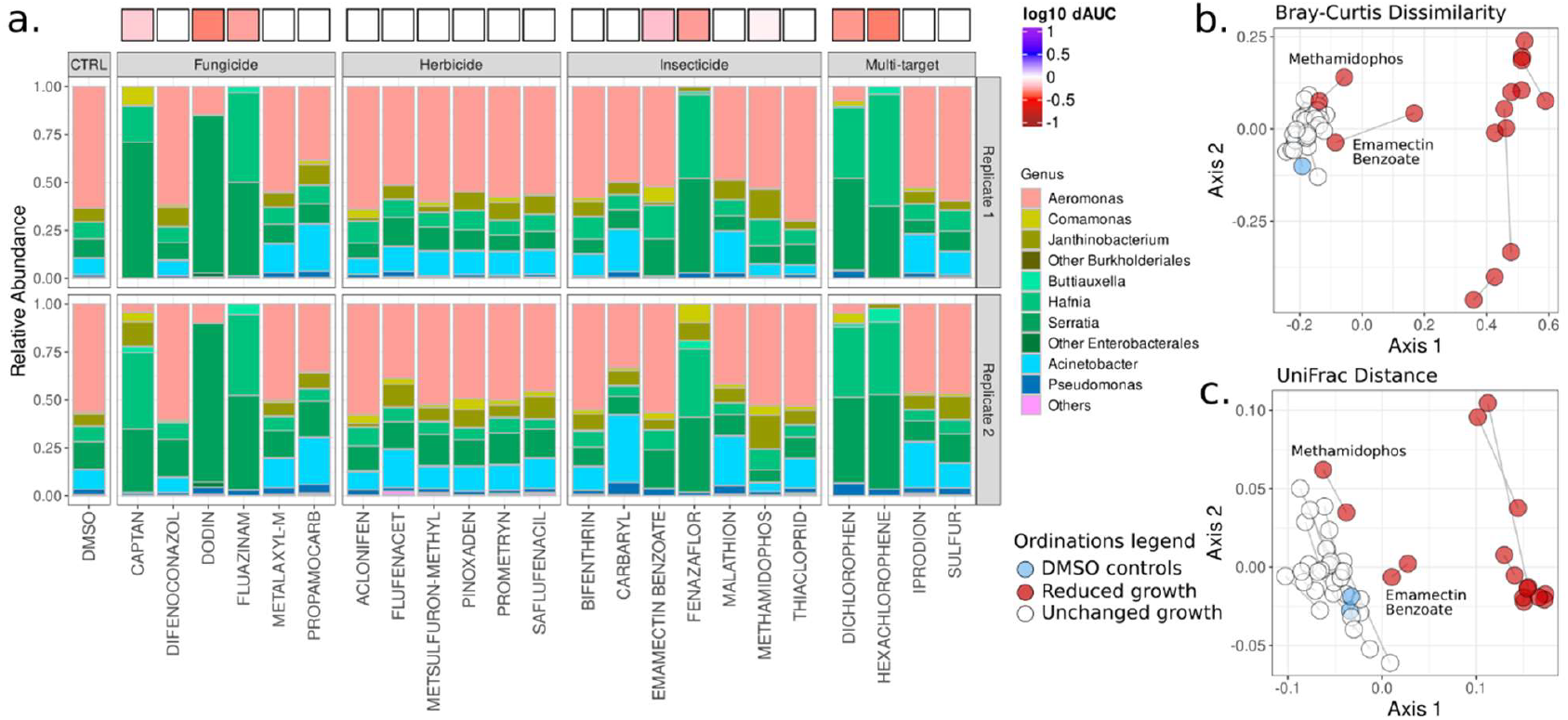
Community composition profiles. **a**. Relative abundance of taxa from Community 9 grown under differing chemical conditions. Growth with DMSO (left panel) is the control condition (CTRL). Relative abundances are aggregated to the genus level, with the most abundant genera shown. Top and bottom panels are sequencing profiles from two replicates for each condition. The colour boxes above the bars show the relative growth (dAUC) on the same log_10_ colour scale as Fig. 2a. Chemicals grouped by broad target classes, with insecticides including acaricides. Multi-target chemicals span more than one of these classifications. **b**. Principal co-ordinates analysis (PCoA) based on Bray-Curtis dissimilarity. The chemicals which don’t impact community functioning (growth) cluster on with the DMSO controls (blue) and represent the chemical treatments which have largely unaffected the community composition. The compositions in the presence of chemicals which impacted community growth (captan, dodin, dichlorophen, emmamectin benzoate, fenazaflor, fluazinam, hexachlorophene and methamidophos; coloured red) are shifted in the ordination space by differing degrees. **c**. PCoA based on weighted UniFrac distance, a measure of phylogenetic dissimilarity of communities. The chemicals which caused reductions in community growth also caused those communities to be more phylogenetically dissimilar to the control condition. In both ordinations (**b**. and **c**.), lines join the two replicate points for each condition. Methamidaphos and emamectin benzoate are labelled to show that these communities cluster near the controls despite showing (moderate) responses in their growth.

We further examined differences in community composition via principle coordinates analysis (PCoA), using the Bray-Curtis dissimilarity metric (Fig. 3b) and using weighted UniFrac distance, a measure of phylogenetic dissimilarity (Fig. 3c). In both cases, we found that replicates generally clustered together, showing repeatability of community assembly in response to the chemical treatments. We found that communities grown in the presence of chemicals which did not affect growth clustered together with the DMSO controls in both ordinations, implying no difference in composition compared to the control condition. We found that the compositions in the presence of chemicals which impacted growth were shifted away from the DMSO controls, in a similar direction in the ordination space, implying a constrained community response to chemical pollutants (Fig. 3b & c). As observed in their taxonomic profiles (Fig. 3a) emamectin benzoate and methamidophos did not induce the same compositional changes as the other impactful chemicals, clustering more with the controls and unimpacted communities than the communities with reduced growth (Fig. 3b & c).

To further test how community composition was impacted by the chemical treatments and to link this to the functional measurements, we quantified three measures of community diversity – species richness, the Shannon index, and the net relatedness index (NRI). The species richness is simply the count of unique species in the communities. The Shannon index measures the diversity based on the species richness but also accounts for the evenness of the community (how evenly the individuals are distributed among species). The NRI quantifies the degree to which species in a community are phylogenetically related, with higher values indicating that species are more closely related than expected by chance. We took the mean of 2 replicates when calculating each of these metrics. Each of these metrics explained the differences in growth of the communities compared to the control to different degrees, with NRI providing the best model fit (linear models: species richness r^2^ = 54%, Shannon index r^2^ = 65%, NRI r^2^ = 77%; Fig. 4). These results show that communities with lower growth compared to the DMSO controls had lower diversity as quantified by the species richness and Shannon index (Fig. 4). They also show that communities with lowered growth were more phylogenetically clustered (positive NRI), i.e. species in these communities were more closely related than in the communities showing no growth response.

**Figure 4:**
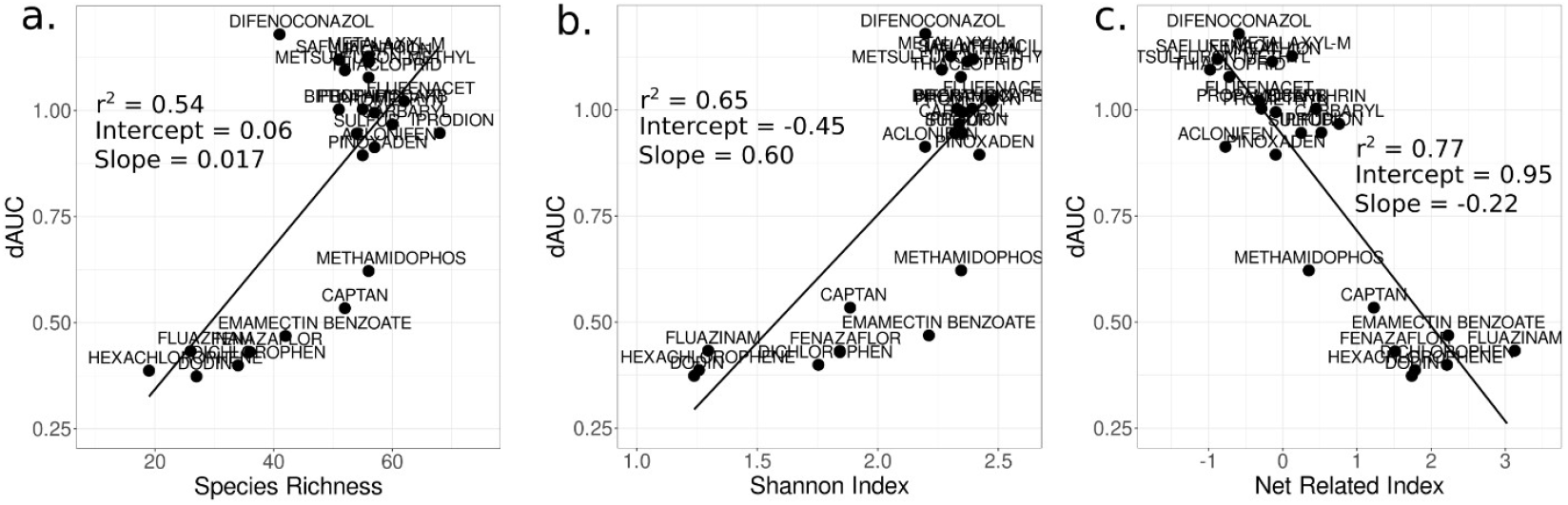
Community diversity, structure and functioning. Here we test the impact of community diversity (species richness, **a**.; Shannon index, **b**.; Net Related Index, **c**.) on functioning (growth). We find that while all three metrics are significant predictors of differences in community growth, the net related index (NRI) is the strongest predictor (r^2^ = 0.77).

The responses of single isolates do not account for biotic interactions between taxa which occur in communities of microbes. We therefore compared the effects of chemicals on specific taxa within a community context to their responses to those chemicals in monoculture. Between our monocultured isolates and sequenced community, there were five shared genera: *Aeromonas, Carnobacterium, Flavobacterium, Pseudomonas* and *Serratia*. We tested whether the functional responses of members of a genus in monoculture were correlated with changes in the abundance of the same genus in a community by comparing the mean functional response (dAUC) of that genus to its aggregated fold change in relative abundance within the community in response to each of the 23 chemical pollutants (Fig. 5). We do not suggest direct causation or dependence between these variables nor assume that one predicts the other, but quantified the strength of relationship between them. We quantified both Pearson and Spearman’s rank correlations (Table 1), capturing linear and monotonic trends respectively. We found significant positive relationships between these chemical response measures for three of our genera: *Aeromonas, Carnobacterium*, and *Flavobacterium*. Conversely, we found a negative relationship for *Serratia*, i.e., *Serratia* relative abundance tended to be increased in the presence of chemicals that caused a decrease in their monoculture growth. However, this negative relationship was not significant by Spearman’s rank after accounting for multiple testing (Table 1). We found no significant relationship between these measures for *Pseudomonas*.

**Table 1:**
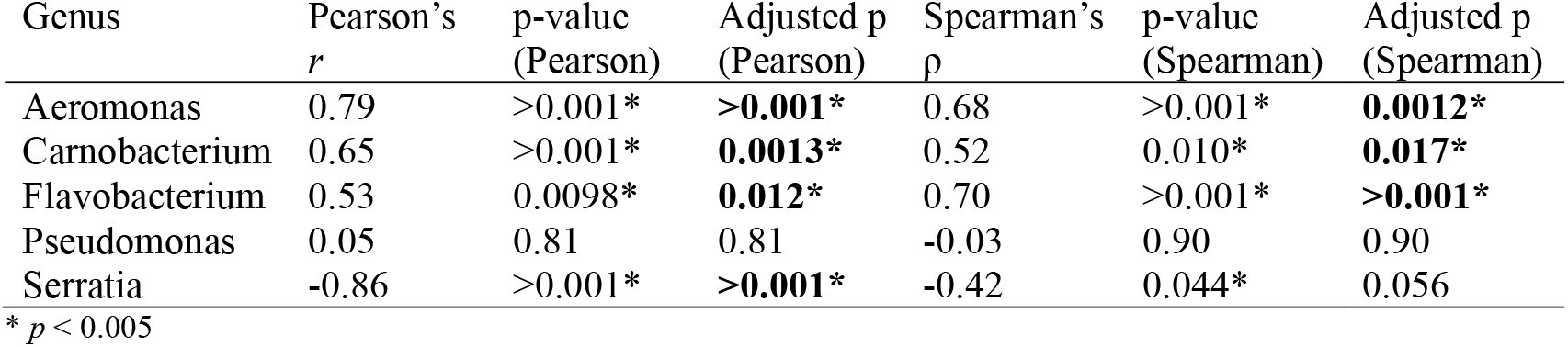
Correlations between monoculture and community responses. Here we provide the Pearson’s and Spearman’s rank test statistics and p-values from correlation tests of isolate growth responses (change in dAUC) against community growth responses (fold change in relative abundance) in the presence of the chemical pollutants (see Fig. 5). The adjusted p-values are after <u>application of a Benjamini-Hochberg false discovery rate adjustment to account for multiple testing.</u>

**Figure 5:**
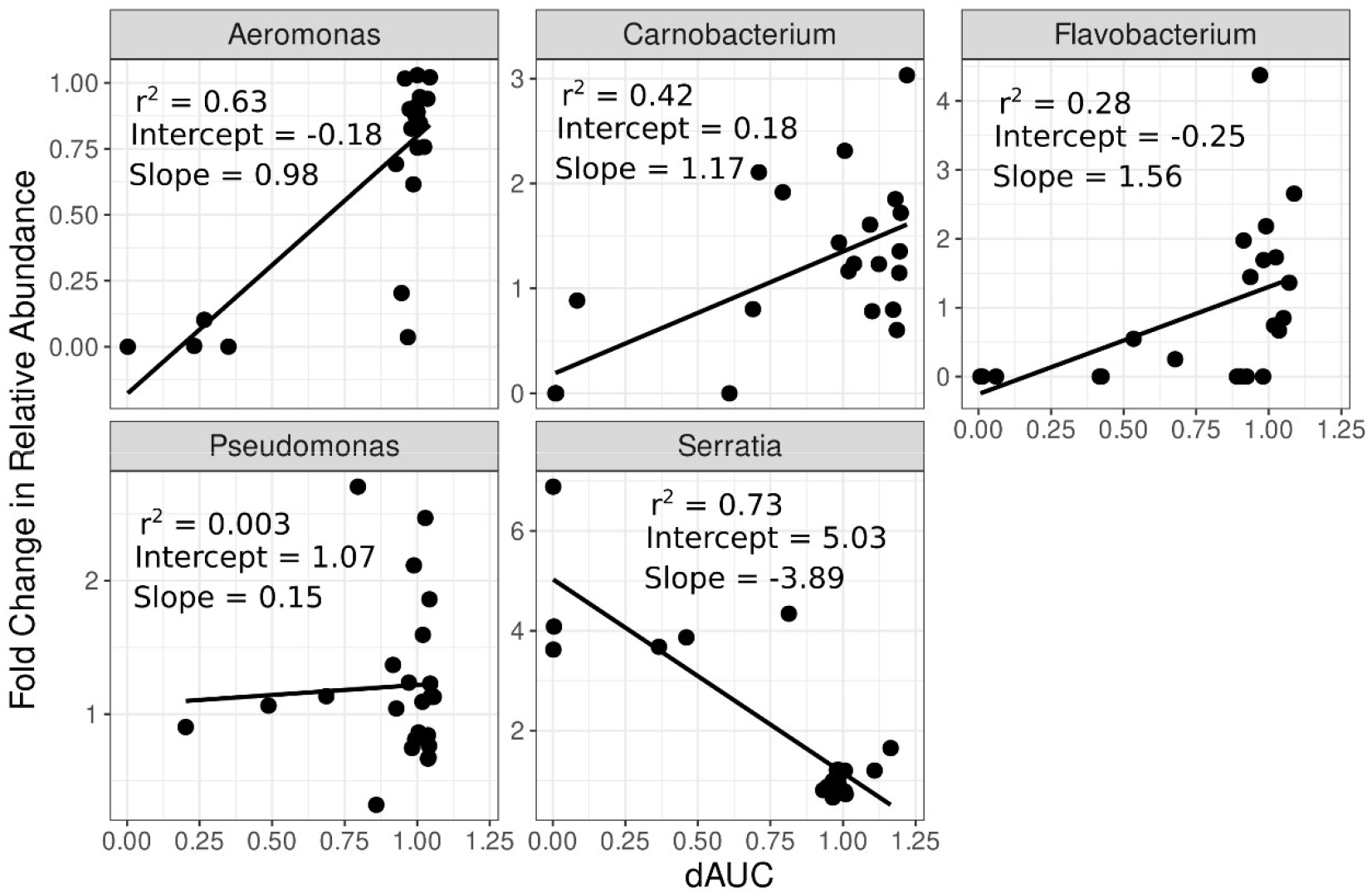
Comparison of shared genera in monoculture and in a community. Here we plot the mean change in growth compared to control growth (dAUC) for five focal genera in the presence of each chemical pollutant, against the fold change in abundance of those genera in the community data. For visualisation, here we plot and provide the statistical outputs from linear regressions, for comparison to the correlation tests in Table 1.

## Discussion

Our study subjected bacterial isolates and whole bacterial communities to a large collection of environmental pollutants and showed the predictability of microbial responses to those chemical pollutants. We found phylogenetic signal in the growth responses of bacterial isolates to chemicals, that is, evolutionarily similar taxa responded similarly to the same chemical treatments. This means that we can use the responses of target taxa to infer the responses of other, similar species. We also found nestedness in the responses of isolates, with some chemicals impacting all or most of the isolates, but others impacting smaller subsets of the bacteria. This shows that within our dataset there were resilient taxa which were impacted by only a small number of chemicals, and more sensitive taxa that were impacted by many more of the chemicals. Many of the broad-spectrum chemicals here also showed broad-spectrum activity in screens of human gut bacteria, further highlighting the generalisability of their off-target activity (Lindell et al., 2024).

The single strain results were reflected in the community analysis. The chemicals which most impacted the isolates also most impacted the community that we sequenced; reducing both the community’s growth and diversity. A key finding is that the phylogenetic structure of the community was also altered in the microcosms with lower growth. We quantified the net relatedness index (NRI) which gives an overall view of the community’s phylogenetic structure, providing insight into broad ecological and evolutionary processes, such as large-scale environmental filtering (Webb et al., 2002). Negative values of NRI imply phylogenetic overdispersion (species in a community are more distantly related than expected by chance), whereas positive values imply phylogenetic clustering (species are more closely related than expected) (Cooper et al., 2008; Webb et al., 2002). Alternatively, community assembly may not be strongly influenced by phylogeny (Cooper et al., 2008), generally -2 < NRI < 2 implies stochastic community assembly, with NRI < - 2 or NRI >2 implying strong overdispersion or clustering respectively. NRI can therefore detect patterns where environmental conditions select for closely related species, leading to phylogenetic clustering (Cavender-Bares et al., 2009; Webb et al., 2002). Our results indicate a shift in community structure from primarily stochastic community assembly under control conditions to strong environmental selection in the communities with lowered growth. This is expected if clades of closely related taxa are lost from the community, as predicted from the phylogenetic signal in the responses of isolates. We also quantified UniFrac distance, the pairwise phylogenetic dissimilarities between the sequenced communities. We found that the communities showing no change in growth tended to cluster together and with the controls in ordination space based on UniFrac distance, while the communities whose growth was impacted clustered away from the controls. This further highlights the use of phylogenetic measures for diagnosing disturbed microbial communities.

None of the isolates tested showed a response to just one chemical, more often they showed similar responses to many of the chemical pollutants. Similarly, few chemicals impacted only one of the isolates tested. From a biosensors perspective, it may therefore be difficult to pinpoint contamination by a specific chemical pollutant based on the response of single isolate “biosensors”. Indeed, previous work has identified groups of taxa which may be indicative of multiple environmental variables (Sperlea et al., 2021). We also found a lack of sensitivity in our community responses – many chemicals which impacted the communities caused similar shifts in the community composition. This suggests the need for a broader approach when using microbial communities as bioindicators. Our findings of sets of sensitive and resilient taxa in both our isolate and community work suggest that loss of sensitive taxa from a community may be an early indication of environmental perturbation. However, we also found that the sensitivities of certain taxa in a community context couldn’t always be inferred from their monoculture responses. Indeed, we found increased relative abundance of *Serratia* in communities dosed with the same chemicals that elicited a negative response in our *Serratia* isolate when grown in monoculture. This may be due to the compositional nature of microbial community datasets, which are difficult to interpret in terms of changes in abundances of certain taxa in a community, when we can only obtain specimens of a certain read depth (Weiss et al., 2017). Furthermore, the biological responses of a taxon may simply be different in a community context where dynamics are influenced by ecological interactions not reflected in monoculture growth (Gralka et al., 2020). This also raises questions, as in previous work (Smith et al., 2024), about the usage of model organisms in ecotoxicology testing, given the difficulties in generalising monoculture responses to broader contexts. Our phylogenetics results show that instead of focusing on the direct responses of indicator taxa, we can infer the loss of sensitive taxa from increased phylogenetic clustering of microbial communities, suggesting that a simple metric (NRI) could be used to move forward with microbial-based bio-monitoring.

A key challenge in using microbial community data for routine water quality monitoring is the time and cost involved with sequencing approaches. Traditional sequencing technologies, such as Illumina, are resource-intensive and can take days to generate results. However, recent advancements in sequencing technology, particularly the use of portable Nanopore sequencing devices, offer a promising alternative. Indeed, studies have already demonstrated the feasibility of using Nanopore technology in the field for rapid sequencing analyses (Menegon et al., 2017; Parker et al., 2017; Tyler et al., 2023). The ability to monitor ecosystems in real time could revolutionise bioindication practices, providing a more immediate understanding of pollutant impacts. We highlight here that we generated the sequencing data for the current study using a portable Nanopore MinION device, further showing the utility of this technology.

Despite the potential for biomonitoring, it is important to note that our work was conducted in a controlled lab environment, growing both isolates and microbial communities in culture media. Therefore, while our findings show promise for future applications, they need to be validated in the field where microbial communities are influenced by a range of variables not present in the lab, such as natural variations in nutrient levels, temperature, and flow conditions. Furthermore, real-world environments often involve exposure to mixtures of pollutants rather than single compounds. This adds a layer of complexity, as it remains unclear whether community responses to combined stressors are as predictable and modellable as those to individual pollutants (Morris et al., 2022; Orr et al., 2020; Smith et al., 2024; Turschwell et al., 2022). Future research should investigate these dynamics, particularly to explore whether responses to mixtures follow the same patterns of nestedness and phylogenetic signal we observed with single chemicals.

More generally, our results have importance for understanding the responses of microbial communities to perturbations. Our finding of broad repeatability in the community composition in response to different chemicals, while useful for biomonitoring processes, is also informative for understanding microbial community assembly. We find that community assembly may have different end-points depending upon the chemical perturbation, but that replicates converge on the same composition. This is in agreement with recent work showing that microbial community assembly converges on a small number of set structural compositions (Pascual-garcía et al., 2023). Furthermore, our data suggest that microbial communities exhibit some degree of resilience, with communities showing greater capacity to tolerate chemical stress than individual bacterial isolates. For example, while hexachlorophene severely inhibited the growth of most isolates, all tested communities maintained some growth in its presence, highlighting the importance of biotic interactions and functional redundancy in determining ecosystem resilience (Biggs et al., 2020). However, this resilience was not without limits. In cases where chemicals led to a clear loss in community diversity, we also observed a corresponding decrease in growth, indicating a potential lack of functional redundancy. This suggests that while communities may be more robust than individual isolates, there is a tipping point where loss of diversity directly translates to loss of ecosystem function. These results highlight the importance of protecting microbial diversity in natural ecosystems, as it underpins both their ecological resilience and biogeochemical functions (Cavicchioli et al., 2019).

Here we move away from a focus on specific taxa, and suggest a broad approach that can be more easily incorporated into environmental monitoring. The development of high-throughput, cost-effective in-situ sequencing technologies, coupled with the phylogenetic approaches we have outlined, has the potential to streamline water quality assessments. Future work should focus on refining these tools for field applications, particularly by testing microbial community responses to a broader range of ecosystems and pollutant mixtures directly in the field. Ultimately, we envision a future where microbial-based bioindicators become an integral part of ecosystem management, enabling real-time monitoring and rapid responses to environmental threats.

## Methods

### Bacterial isolation and culture

In these experiments we assayed 26 strains of bacteria and six whole microbial communities from our -80°C stored culture library. 20 strains of bacteria were isolated from pristine freshwater stream sediments from Iceland; the isolation and identification methods have been described previously (Smith et al., 2024). The six whole microbial communities were obtained from the original sediments that the Icelandic strains were isolated from. Note that the communities are numbered based on their original stream locations, and thus their numeric identifier is not representative of their position in this experiment. The remaining six bacterial isolates were obtained from soils from Silwood Park, Berkshire, UK, which have a history of exposure to agricultural pollution as described previously (Mombrikotb et al., 2022). We selected these strains as representing a wide diversity of taxa in our isolate library for robustness of our phylogenetic analyses (see Fig. 2a), but also, crucially, for their capacity to be reliably revived in a rich culture media and grown to carrying capacity within a short time period (72 hours). Bacterial strains and communities were revived from frozen glycerol stocks and grown to carrying capacity in Luria–Bertani (LB) media (Sigma-Aldrich) before chemical toxicity testing, as ascertained through absorbance readings at 600nm (A_600_).

### Chemical toxicity experiments

A previous study tested off-target toxicity of 1076 industrial and agricultural chemicals on human gut bacteria, and found that 168 were active against at least one bacterial strain (Lindell et al., 2024). Here, we tested the toxicity of those 168 chemicals on our environmental bacteria and communities. Stock chemicals were prepared in DMSO following the same methodology as previously (Lindell et al., 2024). Bacterial cultures were grown to carrying capacity over 4 days, and then inoculated into clear 96-well assay plates via a 1:100 dilution into fresh media adjusted with the chemicals at a concentration of 20µM (1% DMSO). Control wells were prepared with media containing 1% DMSO but no chemical. Further wells with no bacterial inoculant were also prepared as controls for contamination. Growth was assayed by measuring A_600_ once per hour, for 72 hours using an automated plate reader (Synergy 2, Agilent BioTek) and stacker (Biostack). Plates were briefly shaken before reading to homogenise samples and disrupt biofilm formation. Three independent biological replicates of each bacterial-chemical combination were carried out on different days and in different plate layouts of the chemicals to prevent edge effects. For community assays, plates were subsequently stored at -80°C for DNA extraction.

### Quantification of chemical impacts on growth

A chemical may impact different or multiple metrics of bacterial growth (e.g., the lag time, maximum growth rate, carrying capacity), therefore we calculated the area under the growth curve (AUC) as an overall metric of fitness, as previously (Smith et al., 2024). AUC was calculated by fitting a spline curve to each of the growth curves, then integrating over the 72hr growth period. To determine whether chemicals had a significant impact on AUC for each bacterial culture tested, we performed a Dunnett’s test against the control (DMSO only) wells. For downstream analysis we calculated the ratio of growth in the presence of a chemical pollutant to the growth in the presence of DMSO (control) to give the relative growth:

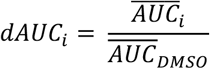

where *i* is one of the 168 chemicals tested and the bar represents the mean of biological replicates. We calculated a dAUC value for each bacterial x chemical combination in this manner.

All growth calculations were performed in R, version 4.3.2 (R core Team, 2022).

### Nanopore sequencing of ‘community 9’

Based on the dAUC results, we selected 23 chemical treatments to sequence the post-growth assay communities for ‘community 9’, as well as the DMSO control (see Supplementary Table S1 for details). We sequenced two replicates of each treatment, for a total of 48 samples sequenced from this community. DNA extraction was performed using a Quick-DNA Fungal/Bacterial Miniprep Kit (Zymo, USA). The Nanopore 16S barcoding kit-24 (SQK-16S024) was used to prepare 16S amplicon libraries for sequencing on a MinION Mk1B device (Oxford Nanopore Technologies, UK), following the manufacturer’s protocol. Barcoded PCR products were quantified using a Qubit fluorometer and adjusted to 10ng/µl with Tris-HCl for each sample. Barcoded samples were then pooled into two libraries of 24 samples, loaded onto a MinION flow cell (R 9.4.1), and read for 12 hours. The flow cell was washed between libraries using the Nanopore flow cell wash kit (EXP-WSH004).

The Nanopore runs of the two libraries generated a total of 3.24 million and 2.68 million reads respectively (>100k reads per sample), with an N50 of 1.65kb. Basecalling, barcode trimming, adapter trimming and read splitting were performed in Guppy version 6.4.6, using the high accuracy model for basecalling. Reads were then size filtered to retain sequences of 1300-1950bp in length using SeqKit version 2.30., following (Matsuo et al., 2021). Species level taxonomic abundances were then estimated using Emu, version 3.4.4 (Curry et al., 2022), with reads mapped to the prebuilt 16S database (a combination of rrnDB v5.6 and NCBI 16S RefSeq from 17-09-2020).

### Phylogenetic reconstruction

We constructed a phylogeny for the bacterial isolates in our experiments using their 16S sequences. These are the 16S sequences directly collected from those strains as part of the process to characterise them. Following previous methodology (Smith et al., 2022), these sequences were aligned to the SILVA 16S reference database using the SILVA Incremental Aligner (SINA) (Pruesse et al., 2012). Based on this alignment we inferred 100 phylogenetic trees in RAxML (v8.1.1) using a general time reversible substitution model with gamma distributed rates, and selected the tree with the highest log-likelihood as the consensus phylogeny (Fig. 2a). For the sequenced ‘community 9’ samples we extracted the 16S sequences from the Emu database matching every unique taxon identified across the sequencing runs. We then performed the same process in SILVA and RAxML to produce a phylogeny representing the community (Supplementary Fig. S3).

### Statistical Analysis

To test for phylogenetic signal in the responses of bacterial isolates to the suite of chemical pollutants, we used the dAUC as the phenotype for each bacteria-chemical combination. We calculated the pairwise Euclidean distance of phenotypic responses between the different strains and compared the pairwise distance matrix to the pairwise distances extracted from the phylogeny using a Mantel test (Fig. 2b). As previously, we used Kendall’s rank correlation τ as the test statistic for the Mantel test due to the non-parametric distribution of the pair-wise phenotypic distances (Smith et al., 2024). The Mantel test was performed using the ‘*Vegan’* R package (v2.6-8) (Oksanen et al., 2022).

To analyse microbial community dissimilarities, we calculated the Bray-Curtis dissimilarities based on the taxonomic abundance profiles in ‘*Vegan’*. We used the previously constructed community phylogeny to calculate UniFrac distances weighted by the taxon abundances, using the *‘phyloseq’* R package (v1.46) (McMurdie & Holmes, 2013). We visualised these as ordinations using principal co-ordinate analysis (PCoA) implemented in the *‘ape’* R package (v5.8) (Paradis & Schliep, 2019). For the community data we also quantified measures of community diversity – species richness and Shannon diversity – using the *‘Vegan’* package. We also calculated the mean phylogenetic distance (MPD) between taxa within each sequenced community sample using the *‘pez’* R package (v1.2-4) (Pearse et al., 2015). We converted this to the net related index (NRI) as the negative of the MPD.

## Acknowledgements

We would like to thank Kiran Patil and Stephan Kamrad for providing stock solutions of the chemical library and for helpful discussions around the experimental methodology. We thank Aiden Zhang for help performing growth assays.

This work was funded by a Natural Environment Research Council grant NE/S000348/1.

## References

Amend, A. S., Martiny, A. C., Allison, S. D., Berlemont, R., Goulden, M. L., Lu, Y., Treseder, K. K., Weihe, C., & Martiny, J. B. H. (2016). Microbial response to simulated global change is phylogenetically conserved and linked with functional potential. ISME Journal, 10(1), 109–118. 10.1038/ismej.2015.96

Astudillo-García, C., Hermans, S. M., Stevenson, B., Buckley, H. L., & Lear, G. (2019). Microbial assemblages and bioindicators as proxies for ecosystem health status: potential and limitations. Applied Microbiology and Biotechnology, 103(16), 6407–6421. 10.1007/s00253-019-09963-0

Bani, A., Randall, K. C., Clark, D. R., Gregson, B. H., Henderson, D. K., Losty, E. C., & Ferguson, R. M. W. (2022). Mind the gaps: What do we know about how multiple chemical stressors impact freshwater aquatic microbiomes? In Functional Microbiomes (1st ed., Vol. 67). Elsevier Ltd. 10.1016/bs.aecr.2022.09.003

Biggs, C. R., Yeager, L. A., Bolser, D. G., Bonsell, C., Dichiera, A. M., Hou, Z., Keyser, S. R., Khursigara, A. J., Lu, K., Muth, A. F., Negrete, B., & Erisman, B. E. (2020). Does functional redundancy affect ecological stability and resilience? A review and meta-analysis. Ecosphere, 11(7). 10.1002/ecs2.3184

Cavender-Bares, J., Kozak, K. H., Fine, P. V. A., & Kembel, S. W. (2009). The merging of community ecology and phylogenetic biology. Ecology Letters, 12(7), 693–715. 10.1111/j.1461-0248.2009.01314.x

Cavicchioli, R., Ripple, W. J., Timmis, K. N., Azam, F., Bakken, L. R., Baylis, M., Behrenfeld, M. J., Boetius, A., Boyd, P. W., Classen, A. T., Crowther, T. W., Danovaro, R., Foreman, C. M., Huisman, J., Hutchins, D. A., Jansson, J. K., Karl, D. M., Koskella, B., Mark Welch, D. B., … Webster, N. S. (2019). Scientists’ warning to humanity: microorganisms and climate change. Nature Reviews Microbiology. 10.1038/s41579-019-0222-5

Cook, L. S. J., Briscoe, A. G., Fonseca, V. G., Boenigk, J., Woodward, G., & Bass, D. (2024). Microbial, holobiont, and Tree of Life eDNA / eRNA for enhanced ecological assessment. Trends in Microbiology, 1–18. 10.1016/j.tim.2024.07.003

Cooper, N., Rodríguez, J., & Purvis, A. (2008). A common tendency for phylogenetic overdispersion in mammalian assemblages. Proceedings of the Royal Society B: Biological Sciences, 275(1646), 2031–2037. 10.1098/rspb.2008.0420

Curry, K. D., Wang, Q., Nute, M. G., Tyshaieva, A., Reeves, E., Soriano, S., Wu, Q., Graeber, E., Finzer, P., Mendling, W., Savidge, T., Villapol, S., Dilthey, A., & Treangen, T. J. (2022). Emu: species-level microbial community profiling of full-length 16S rRNA Oxford Nanopore sequencing data. Nature Methods, 19(7), 845–853. 10.1038/s41592-022-01520-4

Gainsbury, A. M., & Colli, G. R. (2019). Phylogenetic community structure as an ecological indicator of anthropogenic disturbance for endemic lizards in a biodiversity hotspot. Ecological Indicators, 103(February), 766–773. 10.1016/j.ecolind.2019.03.008

Gralka, M., Szabo, R., Stocker, R., & Cordero, O. X. (2020). Trophic Interactions and the Drivers of Microbial Community Assembly. Current Biology, 30(19), R1176–R1188. 10.1016/j.cub.2020.08.007

Grossart, H. P., Massana, R., McMahon, K. D., & Walsh, D. A. (2020). Linking metagenomics to aquatic microbial ecology and biogeochemical cycles. Limnology and Oceanography, 65(S1), S2–S20. 10.1002/lno.11382

Helmus, M. R., Keller, W., Paterson, M. J., Yan, N. D., Cannon, C. H., & Rusak, J. A. (2010). Communities contain closely related species during ecosystem disturbance. Ecology Letters, 13(2), 162–174. 10.1111/j.1461-0248.2009.01411.x

Keck, F., Rimet, F., Bouchez, A., & Franc, A. (2016). Phylosignal: An R package to measure, test, and explore the phylogenetic signal. Ecology and Evolution, 6(9), 2774–2780. 10.1002/ece3.2051

Lindell, A. E., Kamrad, S., Roux, I., Krishna, S., Grießhammer, A., & Patil, K. R. (2024). Off-purpose activity of industrial and agricultural chemicals against human gut bacteria. BioRxiv, 1–26.

Matsuo, Y., Komiya, S., Yasumizu, Y., Yasuoka, Y., Mizushima, K., Takagi, T., Kryukov, K., Fukuda, A., Morimoto, Y., Naito, Y., Okada, H., Bono, H., Nakagawa, S., & Hirota, K. (2021). Full-length 16S rRNA gene amplicon analysis of human gut microbiota using MinIONTM nanopore sequencing confers species-level resolution. BMC Microbiology, 21(1), 1–13. 10.1186/s12866-021-02094-5

McMurdie, P. J., & Holmes, S. (2013). Phyloseq: An R Package for Reproducible Interactive Analysis and Graphics of Microbiome Census Data. PLoS ONE, 8(4). 10.1371/journal.pone.0061217

Menegon, M., Cantaloni, C., Rodriguez-Prieto, A., Centomo, C., Abdelfattah, A., Rossato, M., Bernardi, M., Xumerle, L., Loader, S., & Delledonne, M. (2017). On site DNA barcoding by nanopore sequencing. PLoS ONE, 12(10), 1–18. 10.1371/journal.pone.0184741

Mombrikotb, S. B., Van Agtmaal, M., Johnstone, E., Crawley, M. J., Gweon, H. S., Griffiths, R. I., & Bell, T. (2022). The interactions and hierarchical effects of long-term agricultural stressors on soil bacterial communities. Environmental Microbiology Reports, 14(5), 711–718. 10.1111/1758-2229.13106

Morris, O. F., Loewen, C. J. G., Woodward, G., Schäfer, R. B., Piggott, J. J., Vinebrooke, R. D., & Jackson, M. C. (2022). Local stressors mask the effects of warming in freshwater ecosystems. Ecology Letters. 10.1111/ele.14108

Oksanen, J., Simpson, G. L., Blanchet, F. G., Kindt, R., Legendre, P., Minchin, P. R., O’Hara, R. B., Solymos, P., Stevens, M. H. H., Szoecs, E., Wagner, H., Barbour, M., Bedward, M., Bolker, B., Borcard, D., Carvalho, G., Chirico, M., De Caceres, M., Durand, S., … Weedon, J. (2022). vegan: Community Ecology Package (R package version 2. 6-4). https://cran.r-project.org/package=vegan

Orr, J. A., Vinebrooke, R. D., Jackson, M. C., Kroeker, K. J., Kordas, R. L., Mantyka-Pringle, C., van den Brink, P. J., de Laender, F., Stoks, R., Holmstrup, M., Matthaei, C. D., Monk, W. A., Penk, M. R., Leuzinger, S., Schäfer, R. B., & Piggott, J. J. (2020). Towards a unified study of multiple stressors: Divisions and common goals across research disciplines. Proceedings of the Royal Society B: Biological Sciences, 287(1926). 10.1098/rspb.2020.0421

Paradis, E., & Schliep, K. (2019). Ape 5.0: An environment for modern phylogenetics and evolutionary analyses in R. Bioinformatics, 35(3), 526–528. 10.1093/bioinformatics/bty633

Parker, J., Helmstetter, A. J., Devey, Di., Wilkinson, T., & Papadopulos, A. S. T. (2017). Field-based species identification of closely-related plants using real-time nanopore sequencing. Scientific Reports, 7(1), 1–8. 10.1038/s41598-017-08461-5

Pascual-garcía, A., Rivett, D., Jones, M. L., & Bell, T. (2023). Replaying the tape of ecology to domesticate wild microbiota. BioRxiv, 1–13.

Pearse, W. D., Cadotte, M. W., Cavender-Bares, J., Ives, A. R., Tucker, C. M., Walker, S. C., & Helmus, M. R. (2015). Pez: Phylogenetics for the environmental sciences. Bioinformatics, 31(17), 2888–2890. 10.1093/bioinformatics/btv277

Pruesse, E., Peplies, J., & Glöckner, F. O. (2012). SINA: Accurate high-throughput multiple sequence alignment of ribosomal RNA genes. Bioinformatics, 28(14), 1823–1829. 10.1093/bioinformatics/bts252

R core Team. (2022). R: A language and environment for statistical computing. (4.2.1). R Foundation for Statistical Computing. https://www.r-project.org/

Sagova-Mareckova, M., Boenigk, J., Bouchez, A., Cermakova, K., Chonova, T., Cordier, T., Eisendle, U., Elersek, T., Fazi, S., Fleituch, T., Frühe, L., Gajdosova, M., Graupner, N., Haegerbaeumer, A., Kelly, A. M., Kopecky, J., Leese, F., Nõges, P., Orlic, S., … Stoeck, T. (2021). Expanding ecological assessment by integrating microorganisms into routine freshwater biomonitoring. Water Research, 191(December 2020), 116767. 10.1016/j.watres.2020.116767

Smith, T. P., Clegg, T., Ransome, E., Martin-Lilley, T., Rosindell, J., Woodward, G., Pawar, S., & Bell, T. (2024). High-throughput characterization of bacterial responses to complex mixtures of chemical pollutants. Nature Microbiology. 10.1038/s41564-024-01626-9

Smith, T. P., Mombrikotb, S., Ransome, E., Kontopoulos, D.-G., Pawar, S., & Bell, T. (2022). Latent functional diversity may accelerate microbial community responses to temperature fluctuations. ELife, 11, e80867. 10.7554/eLife.80867

Sperlea, T., Kreuder, N., Beisser, D., Hattab, G., Boenigk, J., & Heider, D. (2021). Quantification of the covariation of lake microbiomes and environmental variables using a machine learning-based framework. Molecular Ecology, 30(9), 2131–2144. 10.1111/mec.15872

Stehle, S., & Schulz, R. (2015). Agricultural insecticides threaten surface waters at the global scale. Proceedings of the National Academy of Sciences of the United States of America, 112(18), 5750–5755. 10.1073/pnas.1500232112

Thompson, M. S. A., Bankier, C., Bell, T., Dumbrell, A. J., Gray, C., Ledger, M. E., Lehmann, K., McKew, B. A., Sayer, C. D., Shelley, F., Trimmer, M., Warren, S. L., & Woodward, G. (2016). Gene-to-ecosystem impacts of a catastrophic pesticide spill: testing a multilevel bioassessment approach in a river ecosystem. Freshwater Biology, 61(12), 2037–2050. 10.1111/fwb.12676

Turschwell, M. P., Connolly, S. R., Schafer, R. B., De Laender, F., Campbell, M. D., Mantyka-Pringle, C., Jackson, M. C., Kattwinkel, M., Sievers, M., Ashauer, R., Cote, I. M., Connolly, R. M., van den Brink, P. J., & Brown, C. J. (2022). Interactive effects of multiple stressors vary with consumer interactions, stressor dynamics and magnitude. Ecology Letters. 10.1111/ele.14013

Tyler, A. D., McAllister, J., Stapleton, H., Gauci, P., Antonation, K., Thirkettle-Watts, D., & Corbett, C. R. (2023). Field-based detection of bacteria using nanopore sequencing: Method evaluation for biothreat detection in complex samples. PLoS ONE, 18(11 November), 1–8. 10.1371/journal.pone.0295028

Wang, B., Zheng, X., Zhang, H., Xiao, F., He, Z., & Yan, Q. (2020). Keystone taxa of water microbiome respond to environmental quality and predict water contamination. Environmental Research, 187(132), 109666. 10.1016/j.envres.2020.109666

Webb, C. O., Ackerly, D. D., McPeek, M. A., & Donoghue, M. J. (2002). Phylogenies and community ecology. Annual Review of Ecology and Systematics, 33, 475–505. 10.1146/annurev.ecolsys.33.010802.150448

Weiss, S., Xu, Z. Z., Peddada, S., Amir, A., Bittinger, K., Gonzalez, A., Lozupone, C., Zaneveld, J. R., Vázquez-Baeza, Y., Birmingham, A., Hyde, E. R., & Knight, R. (2017). Normalization and microbial differential abundance strategies depend upon data characteristics. Microbiome, 5(1), 1–18. 10.1186/s40168-017-0237-y

Wijaya, J., Byeon, H., Jung, W., Park, J., & Oh, S. (2023). Machine learning modeling using microbiome data reveal microbial indicator for oil-contaminated groundwater. Journal of Water Process Engineering, 53(March), 103610. 10.1016/j.jwpe.2023.103610

