## Supplementary Information for "Phylogenetic clustering of microbial communities as a biomarker for chemical pollution"

**Supplementary Figures:**

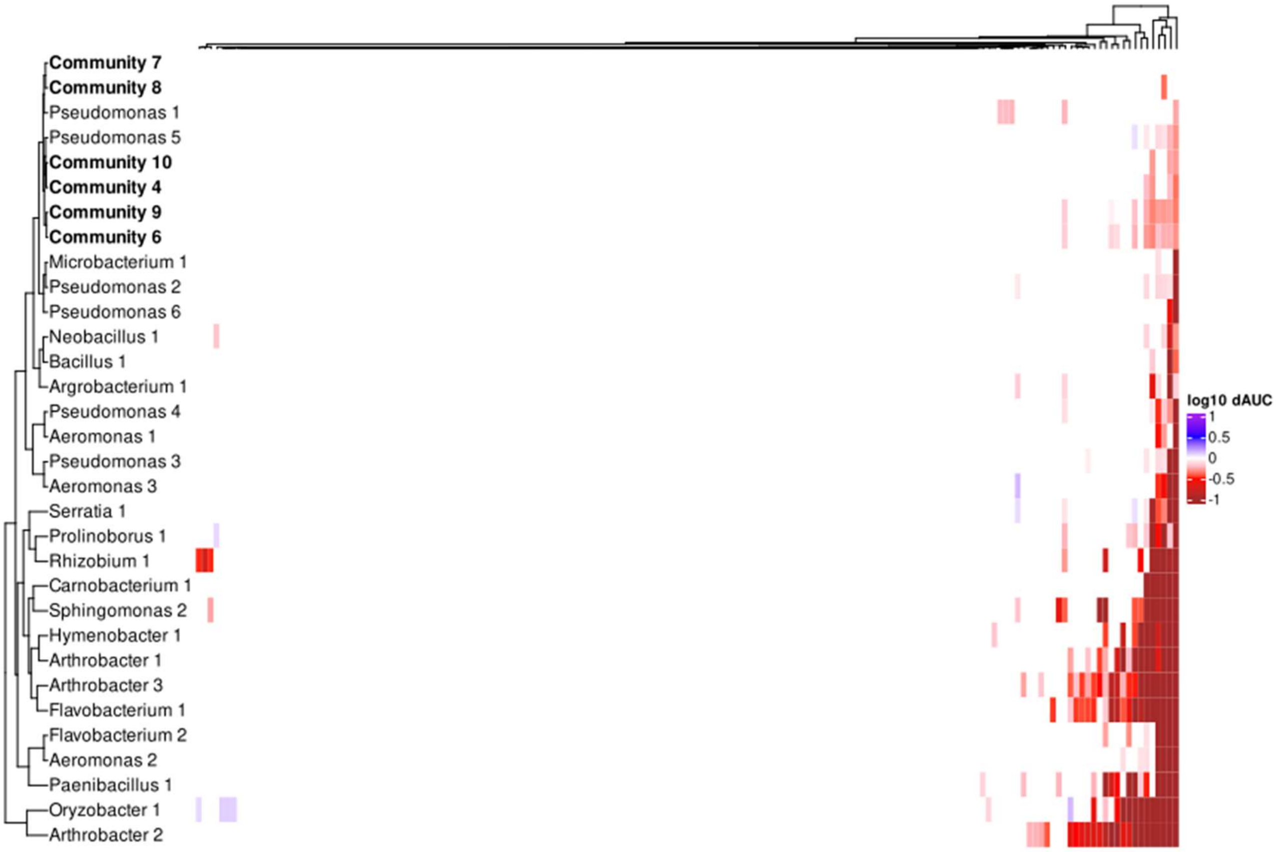


**Figure S1. Heatmap of growth responses to all chemicals.** Most of the 168 chemicals elicit no significant response in either the bacterial isolates or communities tested. Isolates and communities (bold) are clustered by their growth responses. Similar strains cluster together, but not perfectly by phylogeny (see Fig. 1). The communities cluster with the resilient *Pseudomonas* taxa.

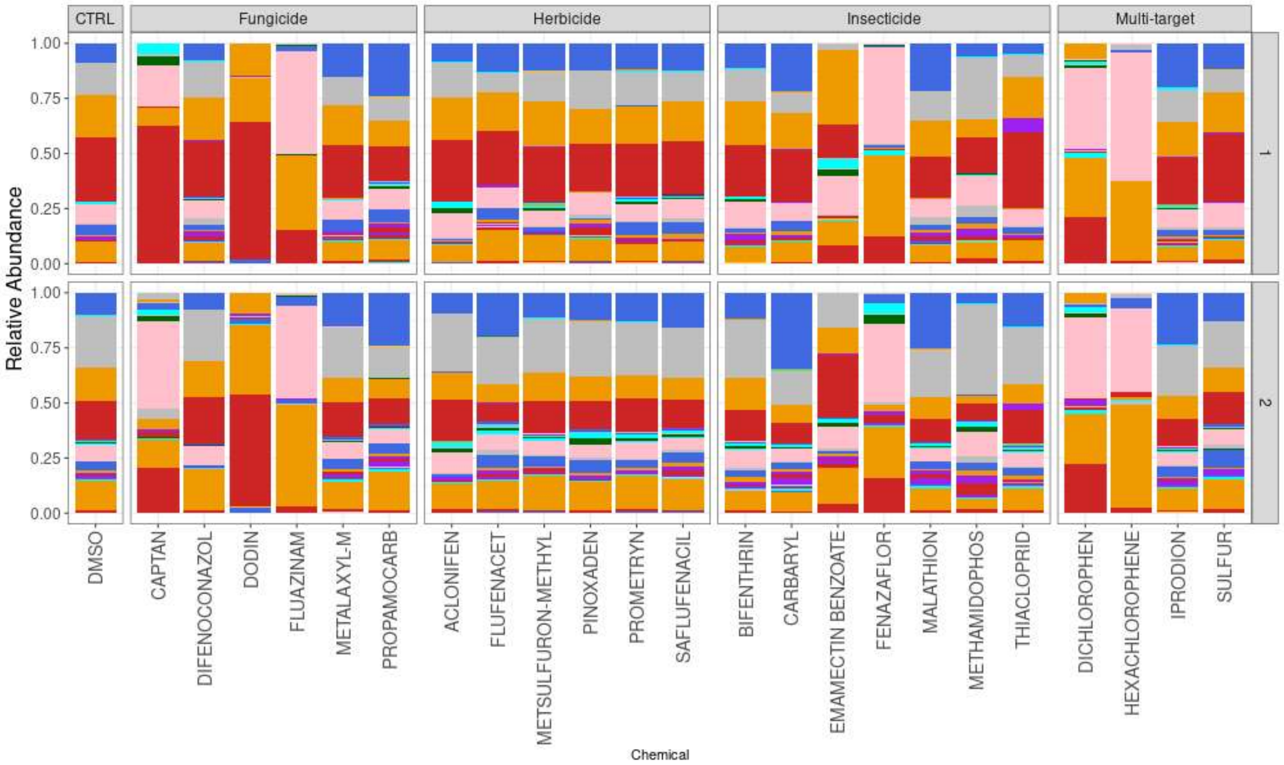


**Figure S2. Species-level community relative abundance profiles.** Here we show the relative abundance profiles from Community 9 using a repeating colour palette to show unique species. We can still see that, as in the aggregated genus-level abundances in Figure 1, the taxonomic profiles in the presence of most chemicals remain very similar to the DMSO control. However, the chemicals which impacted growth show distinct species-level profiles.

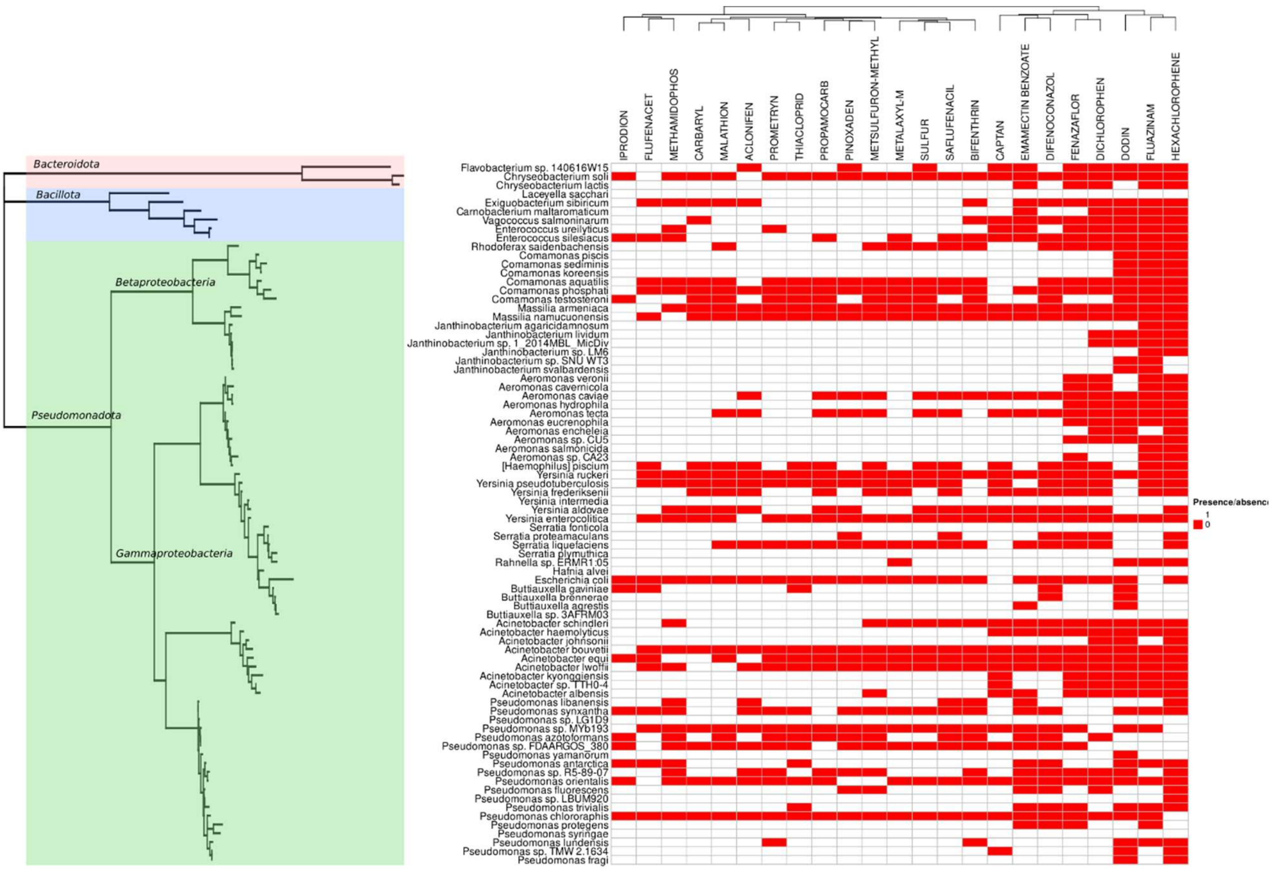


**Figure S3. Phylogeny of species observed in communities grown in the presence of different chemicals.** The phylogeny was built from an alignment of all species observed in our sequencing profiles. Heatmap shows presence of a species (white) or absence (red) when grown in the presence of a given chemical pollutant. Phylogeny is coloured according to the colour scheme of the isolate phylogeny (main text Fig. 2), with main branches of taxa labelled.

**Supplementary Tables:**

**Table S1. Chemicals selected for community compositional analysis.** Here we provide the chemicals and their intended targets in which community 9 was sequenced after growth in the presence of. The dAUC is the relative growth of the community compared to growth in the presence of the DMSO control (mean of 4 replicates). Asterisks denote significantly reduced growth compared to the control (Dunnett's test).

| Chemical | dAUC | Target |
|---|---|---|
| ACLONIFEN | 0.91 | Herbicide |
| BIFENTHRIN | 1.00 | Insecticide |
| CAPTAN | 0.53* | Fungicide |
| CARBARYL | 0.97 | Insecticide |
| DICHLOROPHEN | 0.40* | Multi-target |
| DIFENOCONAZOL | 1.18 | Fungicide |
| DODIN | 0.37* | Fungicide |
| EMAMECTIN BENZOATE | 0.47* | Insecticide |
| FENAZAFLOR | 0.43* | Insecticide |
| FLUAZINAM | 0.43* | Fungicide |
| FLUFENACET | 1.02 | Herbicide |
| HEXACHLOROPHENE | 0.39* | Multi-target |
| IPRODION | 0.95 | Multi-target |
| MALATHION | 1.11 | Insecticide |
| METALAXYL-M | 1.13 | Fungicide |
| METHAMIDOPHOS | 0.62* | Insecticide |
| METSULFURON-METHYL | 1.09 | Herbicide |
| PINOXADEN | 0.89 | Herbicide |
| PROMETRYN | 0.99 | Herbicide |
| PROPAMOCARB | 1.00 | Fungicide |
| SAFLUFENACIL | 1.12 | Herbicide |
| SULFUR | 0.95 | Multi-target |
| THIACLOPRID | 1.08 | Insecticide |
